# Fasting-induced hepatic gluconeogenesis is compromised in *Anxa6^−/−^* mice

**DOI:** 10.1101/2025.01.24.634638

**Authors:** A Alvarez-Guaita, M Bernaus-Esqué, P Blanco-Muñoz, Y Liu, D Sebastian, E Meneses-Salas, MKL Nguyen, A Zorzano, F Tebar, C Enrich, T Grewal, C Rentero

## Abstract

Maintaining constant blood glucose levels is essential for energizing glucose-dependent tissues. During the fed state, insulin lowers elevated blood glucose, while in the fasted state, glucagon maintains blood glucose levels through hepatic stimulation of fatty acid oxidation, glycogenolysis, and gluconeogenesis (GNG). The liver plays a crucial role in these metabolic adaptations. Deregulation of GNG is a hallmark of type 2 diabetes mellitus (T2DM), driven by hepatic insulin resistance, elevated glucagon levels, and excess circulating free fatty acids. Despite normal insulin-sensitive control of glucose levels and effective glycogen mobilization, *Anxa6* knockout (*Anxa6*^−*/*−^) mice display rapid hypoglycaemia during fasting. This metabolic disarrangement, in particular during the early stages of fasting is characterized by a low respiratory exchange ratio (RER) and increased lipid oxidation during the diurnal period, indicating a reliance on lipid oxidation due to hypoglycaemia. Elevated glucagon levels during fasting suggest deficiencies in GNG. Further analysis reveals that *Anxa6*^−*/*−^ mice are unable to utilize alanine for hepatic GNG, highlighting a specific impairment in the glucose-alanine cycle in fasted *Anxa6*^−*/*−^ mice, underscoring the critical role of ANXA6 in maintaining glucose homeostasis under metabolic stress. During fasting, slightly reduced expression levels of alanine aminotransferase 2 (*Gpt2*) and lactate dehydrogenase (*Ldha2*), enzymes converting alanine to pyruvate, and the hepatic alanine transporter SNAT4 might contribute to these observations in the *Anxa6*^−*/*−^ mice. These findings identify that ANXA6 deficiency causes an inability to maintain glycolytic metabolism under fasting conditions due to impaired alanine-dependent GNG.

## BACKGROUND

Circulating blood glucose levels need to be steadily maintained irrespective of dietary nutritional inputs or during fasting periods to supply glucose-dependent tissues (central nervous system, red blood cells and renal medulla). During the fed state, high levels of glucose induce insulin secretion, which stimulates glucose uptake by cells, and activates anabolic pathways such as glycogen production and lipogenesis and inhibits catabolic pathways. In the fasted state, blood glucose levels are maintained by elevated glucagon levels stimulating hepatic release of glucose generated by fatty acid oxidation, glycogenolysis and gluconeogenesis (GNG) [1]. The liver plays an essential role in this metabolic adaptation through housing processes such as lipogenesis, glycogen production, fatty acid oxidation, glycogenolysis and GNG to control glucose homeostasis.

During metabolic stress such as fasting, the liver is responsible for maintaining blood glucose levels [2]. Initially, elevation of blood glucagon levels triggers glucose production and secretion from hepatic glycogenolysis (glycogen breakdown). Once glycogen stores are depleted, glucose is then *de novo* synthetized from various gluconeogenic precursors, including amino acids from extrahepatic tissues. When in excess, dietary amino acids even stimulate hepatic GNG in the fed state [3, 4]. Importantly, deregulation of GNG is a hallmark of type 2 diabetes mellitus (T2DM), and greatly contributes to highly elevated blood glucose levels in diabetic patients. In the latter, GNG is increased due to insulin resistance failing to inhibit hepatic glucose production, elevated glucagon levels activating pro-GNG signalling pathways, and excess of circulating free fatty acids driving GNG [5].

Supply for primary carbon skeletons used for GNG may derive from pyruvate, lactate, glycerol and amino acids. During fasting, glutamine and alanine account for 60-80% of amino acids released from skeletal muscle [6], with alanine being the main hepatic GNG substrate via the Cahill cycle [7, 8]. In hepatocytes, alanine is mainly taken up by the system A ubiquitously expressed Na^+^-coupled neutral amino acid transporter (SNAT) 2 and the liver-specific SNAT4 [9]. Little is known about the *in vivo* regulation of these SNAT transporters, although feedback mechanisms for SNAT4 expression and sinusoidal plasma membrane localization during liver development and regeneration have been reported [10, 11]. Interestingly, the retromer complex, which orchestrates endocytic recycling and maintains cell surface abundance of nutrient transporters [12], delivers SNAT2 to the plasma membrane upon amino acid withdrawal to prevent its degradation in lysosomes [13]. Importantly, the GTPase RAB7 regulates the recruitment of several retromer subunits to endosomes and cargo recognition [14]. This process is inhibited by the RAB7-GTPase activating protein (GAP) TBC1D5 [15], which promotes RAB7 inactivation and enables fusion of autophagosomes to lysosomes [16]. As retromer expression is elevated upon amino acid starvation, this indicates that transcriptional control of endosomal recycling provides an important means for the adaptive regulation of nutrient uptake [13].

Annexin A6 (ANXA6) is a calcium-dependent phospholipid-binding protein highly abundant in the liver that is involved in membrane trafficking [17], epidermal growth factor receptor and RAS/mitogen-activated protein kinase signalling [17, 18], plasma-membrane microdomain organisation [19], and cholesterol homeostasis through the regulation of RAB7 activity [20-22]. Specifically, ANXA6 recruits the RAB7-GAP TBC1D15 to late endosomes, promoting RAB7 inactivation [22]. *Anxa6* knockout (*Anxa6*^−*/*−^) mice appear normal, with slight alterations in cardiomyocyte function [23] and adiponectin secretion [24]. Yet, when assessed under challenging physiological conditions, such as high-fat diet, ANXA6 deficiency identified an inability to inhibit insulin-dependent GNG [25]. Moreover, our recent work revealed ANXA6 to critically control the survival of mice during liver regeneration [10]. During this process, the remnant liver needs to maintain hepatic functions that control blood glucose homeostasis. However, during the hepatic regeneration program, *Anxa6*^−*/*−^ mice were unable to produce glucose *de novo* due to SNAT4 mislocalization, which severely impaired hepatic alanine uptake, the main gluconeogenic substrate during the regeneration process [10].

Little is known about the function of ANXA6 in the regulation of glucose homeostasis under physiological stress conditions such as fasting. Here, we demonstrate that *Anxa6*^−*/*−^ mice showed no differences in blood glucose secretion or absorption capability, yet these mice were unable to maintain blood glucose levels in the initial stages of fasting and during the fasting state. While glycogen mobilization and GNG from pyruvate and glutamine were not affected, *Anxa6*^−*/*−^ mice were unable to produce glucose *de novo* from alanine. After 24 hours of fasting, the liver of *Anxa6*^−*/*−^ mice displayed slightly reduced expression levels of alanine aminotransferase 2 (*Gpt2*), lactate dehydrogenase (*Ldha2*), and the alanine transporter SNAT4 (*Slc38a4*). Altogether our findings identify an unexplored defect upon ANXA6 deficiency related to the feedback control mechanisms that link nutrient sensing with metabolic enzyme expression and acid transporter recycling in the liver of *Anxa6*^−*/*−^ mice.

## MATERIAL AND METHODS

### Animals

Eight-to twelve-week-old C57Bl6/J wild type (WT) and *Anxa6*^−*/*−^ male mice [26] were used for all experiments and were maintained in a 12 h light/dark cycle, allowed food (regular low-cholesterol, low-fat cereale based rodent chow diet (2014 Teklad Global 14% protein rodent maintenance diet; Envigo)) and water *ad libitum*. Every effort was made to minimize animal suffering and to use the minimum number of animals per group and experiment. All the animal care and experimental procedures were approved by the Local Ethical Committee of the University of Barcelona following European (2010/63/UE) and Spanish (RD 53/2013) regulations for the care and use of laboratory animals.

### Indirect calorimetry

Mice energy expenditure was measured by open circuit indirect calorimetry using the Oxymax 8-chamber system (Columbus Instruments). Before recording the rates of oxygen consumption (VO_2_) and carbon dioxide production (VCO_2_), mice were allowed to adapt to the standard Oxymax chambers for 2 days. VO_2_ and VCO_2_ measured at 22°C for 24 h, for 1.5 min in 20-min intervals for each animal. RER, glucose and lipid oxidation were calculated according to the following equations: RER = VCO_2_/VO_2_; Glucose oxidation [g/min] = 4.55 ∗ VCO_2_ - 3.21 ∗ VO_2_; and Lipid oxidation [g/min] = 1.67 ∗ VO_2_ - 1.67 ∗ VCO_2_ [27]. Mice activity (ambulation) was monitored as events of each mice traversing the cage (counts/hour).

### Glucose metabolism in vivo studies

For glucose and insulin tolerance tests, mice were fasted for 5 hours before intraperitoneal (i.p.) injection of 2 g/kg glucose or 0.75 U/kg insulin. For the determination of GNG, mice were fasted for 24 hours prior to i.p. injection of 2 g/kg sodium pyruvate, glycerol, lactate, L-glutamine or L-alanine. Blood glucose levels were determined from blood obtained by a small incision in the mouse tail using a glucometer (Glucocard G+ meter set, Arkray) at 0, 15, 30, 60, and 120 min after injection.

### Glycogen quantification

To assess liver glycogen content, a 200 mg liver sample was homogenized in 1 ml 30% KOH at 100ºC for 10 min. Following the hydrolysis, the samples were allowed to cool to room temperature, and 2 ml of ethanol was added. The mixture was then incubated for 24 h at −20°C to facilitate glycogen precipitation. After incubation, samples were centrifuged at 2,000 × g for 15 min at 4ºC, and the supernatant was discarded. The resulting pellet was resuspended in 3 ml of 1:2 (v/v) mixture of distilled water and ethanol at 4ºC, and the suspension was subjected to a second centrifugation at 2,000 × g for 15 min at 4ºC. The pellet was then resuspended in 1 ml 5N H_2_SO_4_ and incubated at 100ºC for 2 h to ensure complete hydrolysis of the glycogen to glucose. Finally, the samples were neutralized with 1N NaOH using phenolphthalein (Fluka) as pH indicator. Glucose concentrations, representative of glycogen levels, were quantified using the Glucose assay kit (Sigma) according to manufacturer’s instructions.

### Insulin and glucagon quantification

Insulin (US Mouse Insulin ELISA, Mercodia) and glucagon (Glucagon EIA kit, Sigma) plasma levels were determined by ELISA according to manufacturer’s instructions.

### Beta-hydroxybutyrate (BOH, ketone body) quantification

For the determination of blood ketone bodies, blood was collected by intracardiac punction in BD Blood Collection Microtainer tubes (BD PST™ Lithium heparin/gel). Five μl of plasma sample was analysed with Ketone bodies kit (Sigma) following the manufacturer’s instructions.

### RNA extraction and quantitative Real-Time PCR

Total RNA was prepared from mice liver using RNeasy Lipid Tissue Mini Kit (Qiagen) in accordance with the manufacturer’s protocol. 1 μg RNA from each sample was reverse transcribed using High Capacity cDNA Reverse Transcription Kit (Applied Bioscience). In a final volume of 20 μl real-time PCR Brilliant SYBRGreen QPCR Master Mix (Agilent Technologies), 10 μl of 1:20 diluted cDNA as a template and specific primers (see Table 1) together with a standard PCR amplification protocol (10 min at 95ºC, 45 cycles of 30sec at 95ºC, 15sec at 60ºC and 30sec at 72ºC, 10sec at 95ºC and 60sec at 65ºC) and the LightCycler system (Roche Diagnostics) were used for real-time PCR reaction according to manufacturer’s instructions. Relative gene expression data was analysed following the 2^-ΔΔCt^ method [28].

**Table 1.**
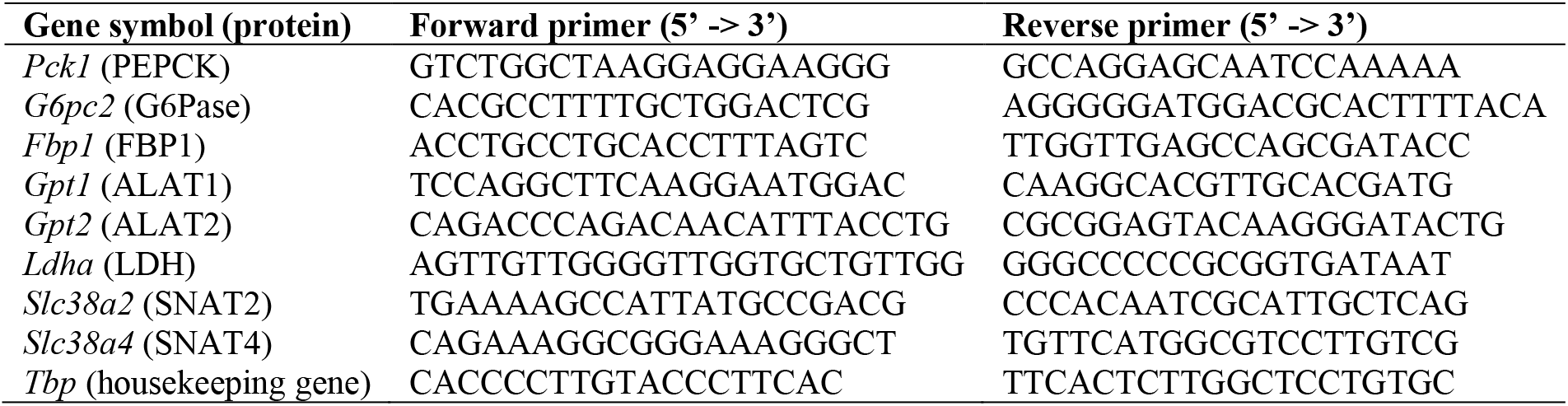
Mouse specific primer sequences for real-time PCR.

### Preparation of liver lysates

All procedures were conducted at 4ºC. Livers were removed from mice and liver tissue samples were placed in Lysing Matrix tubes (MP Biomedicals) with homogenization buffer (10 mM Tris, 150 mM NaCl, 5 mM EDTA, pH 7.5) containing 10 μg/ml of leupeptin and aprotinin, and 1 mM of orthovanadate, NaF and PMSF. Samples were then homogenized in a FastPrep120 homogenizer (MP Biomedicals) and stored at −20ºC.

### Plasma and liver amino acid levels

Plasma amino acid levels were quantified from 100 μl of plasma by cation-exchange chromatography followed by post-column derivatization with ninhydrin and UV detection as described [29]. Proteins were precipitated with 100 μl 10% TCA. NorLeucine served as internal control.

### ALAT activity quantification

For ALAT activity determination, a 200 mg liver sample was homogenized in 1.5 ml of 1M Tris-HCl, 150mM NaCl and 5mM EDTA, and filtered with a 0.45 μm PVDF filter. ALAT activity was then analysed using an autoanalyzer Advia 1650 (Bayer HealthCalre) according to manufacturer’s instructions.

### Western blot analysis

Liver or cell lysates boiled in 1x sample buffer, resolved on SDS-PAGE and transferred to Immobilon-P (Millipore) membranes. Membranes were blocked in 5% non-fat milk, incubated overnight in primary antibodies, washed in TBST, incubated with HRP-conjugated secondary antibodies (see Table 2) and developed using enhanced chemiluminescence ECL (NZYtech) and the density of each spot pixel on the membrane was determined by Image Quant LASS 4000 (GE Healthcare). ImageJ software was used for quantitative analysis of WB bands [30].

**Table 2.** Specific antibodies used in this work.

| Antibody | Company | Application |
| --- | --- | --- |
| anti-SNAT4 (rabbit polyclonal) | Our laboratory [10] | WB (1:1,000) |
| anti-GAPDH (goat monoclonal) | Genescript Cat.No.A00191 | WB (1:10,000) |
| anti-rabbit IgG-HRP (secondary antibody) | Bio-Rad Cat.No.170-6515 | WB (1:3,000) |
| anti-goat IgG-HRP (secondary antibody) | Promega Cat.No.V8051 | WB (1:3,000) |

### Statistics

Data are shown as means ± SEM. Statistical comparison of two groups was performed using a Student’s t test; analysis of interaction was performed with a two-way ANOVA with *ad hoc* Bonferroni post-test. Symbols represent ∗ P<0.05, ∗∗*P*<0.01, ∗∗∗*P*<0.001. Statistical analysis was performed in GraphPad Prism 10.

## RESULTS

### ANXA6 deficiency induces metabolic energetic disarrangement in mice

The liver, together with other organs, coordinates the systemic adaptation to metabolic fluctuations of glucose, fatty acids and amino acid levels. In previous studies we showed that ANXA6 plays an important role in the regulation of lipid and glucose metabolism under high-fat diet [25] or during liver regeneration [10]. However, the potential function of ANXA6 in response to limited dietary nutrient availability remains to be elucidated.

Therefore, we initially assessed the general ability of *Anxa6*^−*/*−^ mice to adapt to physiological changes that require hormone-regulated maintenance of blood glucose levels. We analysed the metabolic response in a normal light-dark cycle, with voluntary and cyclical feeding and fasting periods. WT and *Anxa6*^−*/*−^ mice were scrutinised in metabolic chambers for 24 hours, and oxygen and carbon dioxide levels were monitored every 20 min. The respiratory exchange ratio (RER), reflecting the rates of energy expenditure, and the glucose and lipid oxidation ratios were calculated (Figure 1A-C, Figure S1B-D). WT mice showed RER values around 1 during the last diurnal and the night (active) periods, indicative of carbohydrates being the predominant fuel source (Figure 1A). Interestingly, during the early hours of the diurnal period (6am - 12 noon), WT mice showed lower RER indicating fat oxidation as predominant fuel source until GNG induction in the late light period (12 noon - 6pm). The mean glucose and lipid oxidation levels during these 6-hour periods confirmed lower glucose oxidation during the early diurnal period (6am - 12 noon) (Figure 1B), which was complemented with lipid oxidation as the predominant energetic metabolic pathway (Figure 1C). In contrast, *Anxa6*^−*/*−^ mice exhibited low RER during the whole diurnal (inactive, voluntary fasting) period (6am - 6pm). During this period, this was associated with low glucose oxidation and significantly higher lipid oxidation (Figure 1B-C), reflecting the activation of lipid-oxidation pathways in *Anxa6*^−*/*−^ mice.

**Figure 1.**
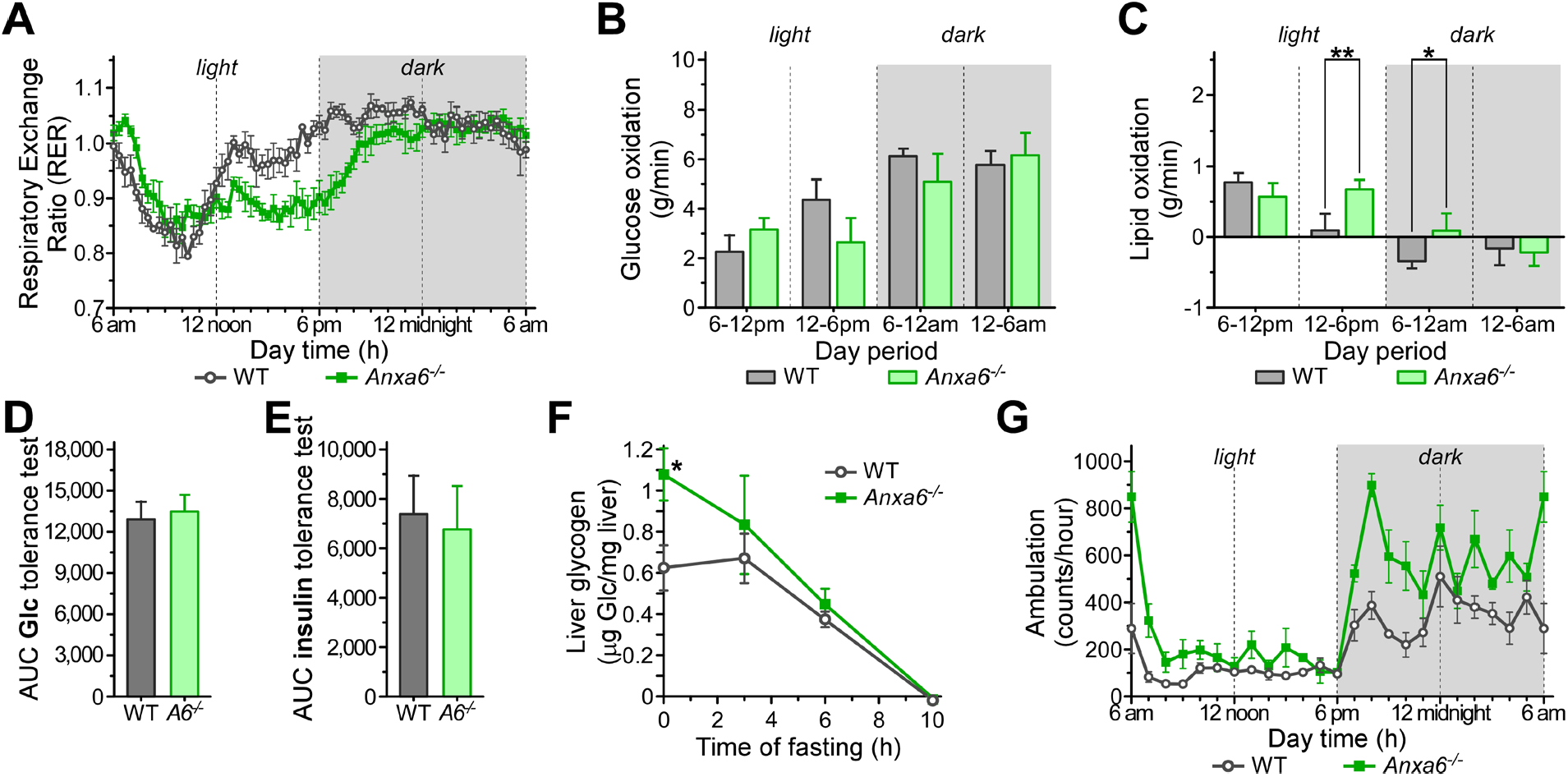
Energetic imbalance during voluntary fasting (light period) in *Anxa6*^−*/*−^ mice. (A) Respiratory exchange ratio (RER) from WT and *Anxa6*^−*/*−^ mice measured every 20 minutes (n=4 mice per group). (B) Glucose oxidation flux expressed as the mean of 6-hour period during day and night-time of WT and *Anxa6*^−*/*−^ mice (n=4 mice per group). (C) Lipid oxidation flux expressed as the mean of 6-hour period during day and night-time of WT and *Anxa6*^−*/*−^ mice (n=4 mice per group). (D) Area under the curve (AUC) from glucose tolerance test of WT and *Anxa6*^−*/*−^ mice (n=6 mice per group) after 5 hours fasting administrating i.p. 2 g/kg glucose. (E) Area under the curve (AUC) from insulin tolerance test of WT and *Anxa6*^−*/*−^ mice (n=6 mice per group) administrating i.p. 0.75 U/kg insulin. (F) Liver glycogen levels in WT and *Anxa6*^−*/*−^ mice (n=3-6 mice per group and time point) during fasting. (G) Mice ambulation per hour from WT and *Anxa6*^−*/*−^ mice (n=4 mice per group).

In line with our previous studies [25], the insulin-sensitive control of systemic glucose levels, glucose absorption and secretion were not negatively affected in the *Anxa6*^−*/*−^ mice during regular feeding conditions (Figure 1D-E). We next analysed the metabolism of glycogen, the initially mobilized hepatic source of glucose during early stages of hypoglycaemia (Figure 1F). As described earlier [10], the amount of hepatic glycogen stored in fed animals was significantly higher in ANXA6-deficient mice. During fasting, *Anxa6*^−*/*−^ more rapidly mobilized their hepatic glycogen stores, involving effective induction of glycogenolysis. However, WT mice increasingly mobilized their hepatic glycogen stores 3 hours after fasting, then reaching similar rates to those observed in the *Anxa6*^−*/*−^ mice (Figure 1F). When body weight, food and water intake, urine volume and defecation were analysed in WT and *Anxa6*^−*/*−^ mice (Figure S2), a statistically significant increase in *Anxa6*^−*/*−^ mice water intake and urine volume was detected. Interestingly, *Anxa6*^−*/*−^ mice showed higher activity both during the light and dark periods, which was statistically significant during the initial phase of these periods (Figure 1G, Figure S1D).

**Figure S1.**
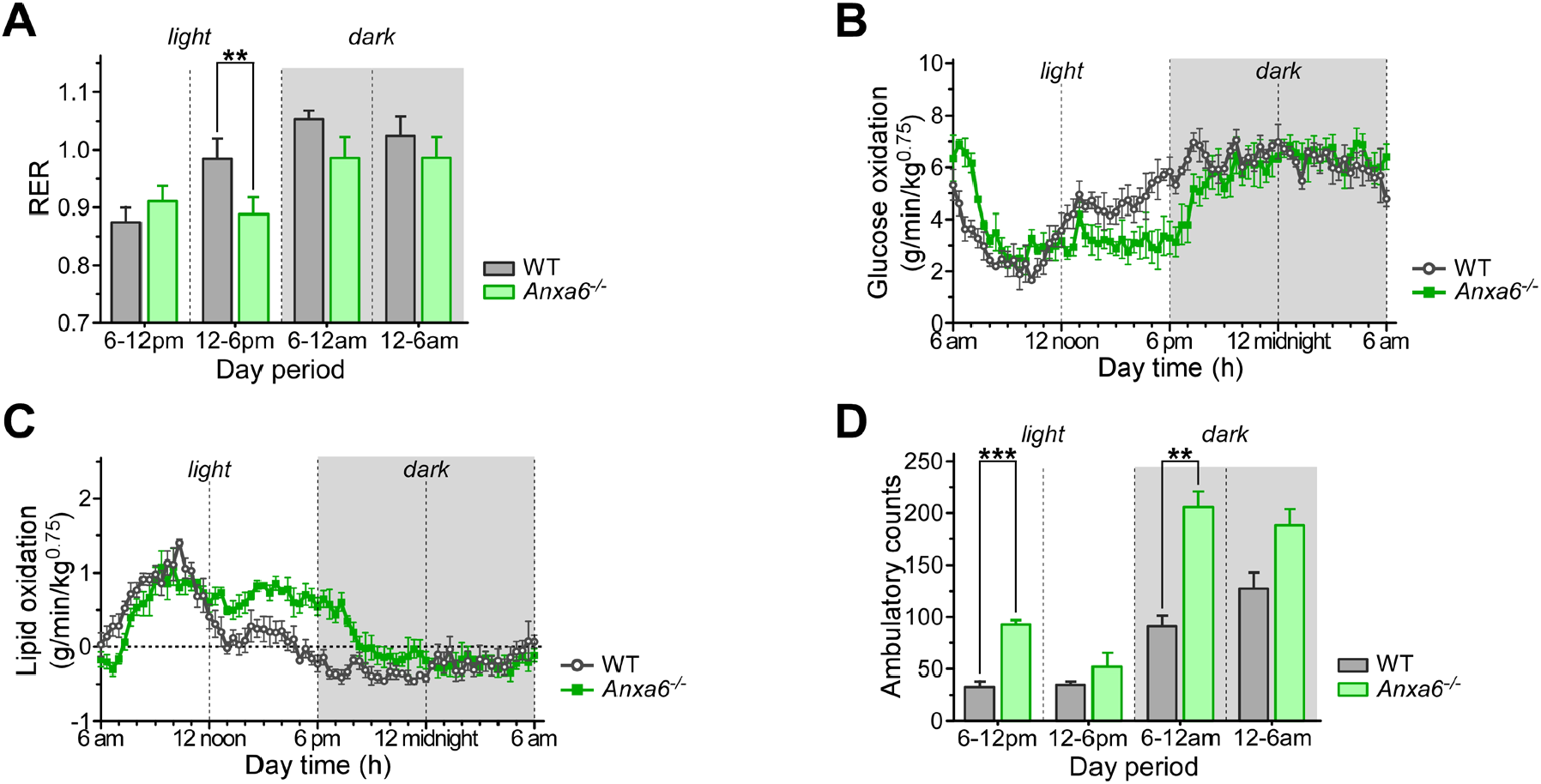
(A) Respiratory exchange ratio (RER) expressed as the mean of 6-hour period during day and night-time of WT and *Anxa6*^−/−^ mice (n=4 mice per group). (B) Glucose oxidation from WT and *Anxa6*^−/−^ mice measured every 20 minutes (n=4 mice per group). (C) Lipid oxidation from WT and *Anxa6*^−/−^ mice measured every 20 minutes (n=4 mice per group). (D) Mice ambulation expressed as the mean of 6-hour period during day and night-time of WT and *Anxa6*^−/−^ mice (n=4 mice per group).

**Figure S2.**
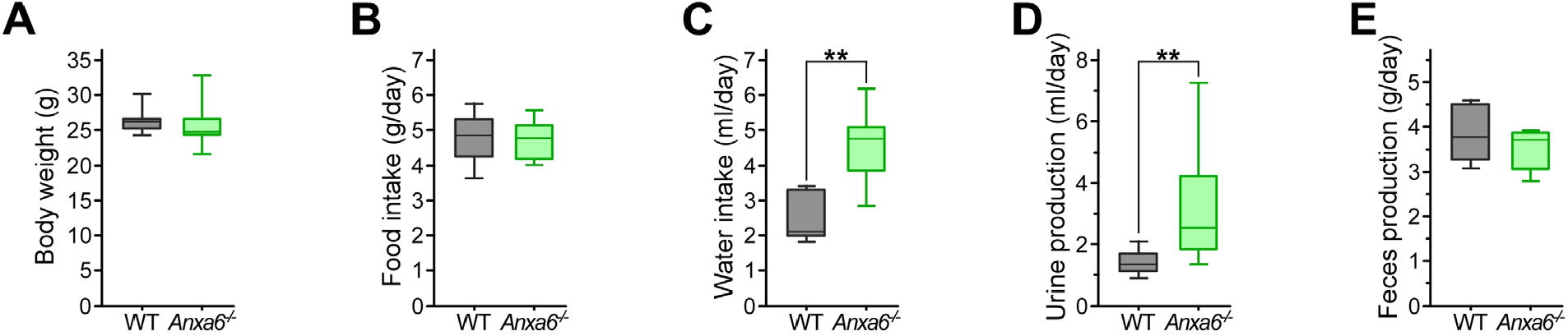
(A) Body weight of 10-week-old WT and *Anxa6*^−/−^ mice (n=7 mice per group). (B) Food intake of WT and *Anxa6*^−/−^ mice (n=7 mice per group). (C) Water intake of WT and *Anxa6*^−/−^ mice (n=7 mice per group). (D) Urine production of WT and *Anxa6*^−/−^ mice (n=7 mice per group). (E) Faeces production of WT and *Anxa6*^−/−^ mice (n=7 mice per group).

Hence, these data suggest energetic alterations of glucose metabolism in *Anxa6*^−*/*−^ mice during the diary light period (voluntary fasting period), which correlates with the recently described GNG impairment during liver regeneration [10]. Yet, ANXA6-deficient mice presented a regular insulin and glucose response and metabolization, although showed higher glycogen stores in the liver.

### Glucose metabolism during fasting in *Anxa6*^−*/*−^ mice

To further characterize the energetic metabolism during fasting, WT and *Anxa6*^−*/*−^ mice were deprived of food and the concentration of circulating blood glucose was measured over 24 hours. Interestingly, and in contrast to the relatively constant glucose levels observed in WT mice during the first 10 hours of fasting, *Anxa6*^−*/*−^ mice displayed a rapid and marked hypoglycaemia already at early fasting time points (Figure 2A). In line with this hypoglycaemic condition, plasma insulin levels rapidly dropped and were significantly lower in *Anxa6*^−*/*−^ mice when compared with their WT counterparts (Figure 2B). On the other hand, plasma glucagon levels were significantly higher during the early fasting period in *Anxa6*^−*/*−^ mice compared to controls (Figure 2C). In spite of this, RER, glucose and lipid oxidation after 12 hours fasting yielded non-significant differences between both mice strains (Figure 2D-F). Also, when lipid oxidation was measured by means of serum ketone bodies (β-hydroxybutyrate), a slight but not significant reduction of serum ketone bodies was evident in *Anxa6*^−*/*−^ mice after 12 hours of fasting (Figure 2G).

**Figure 2.**
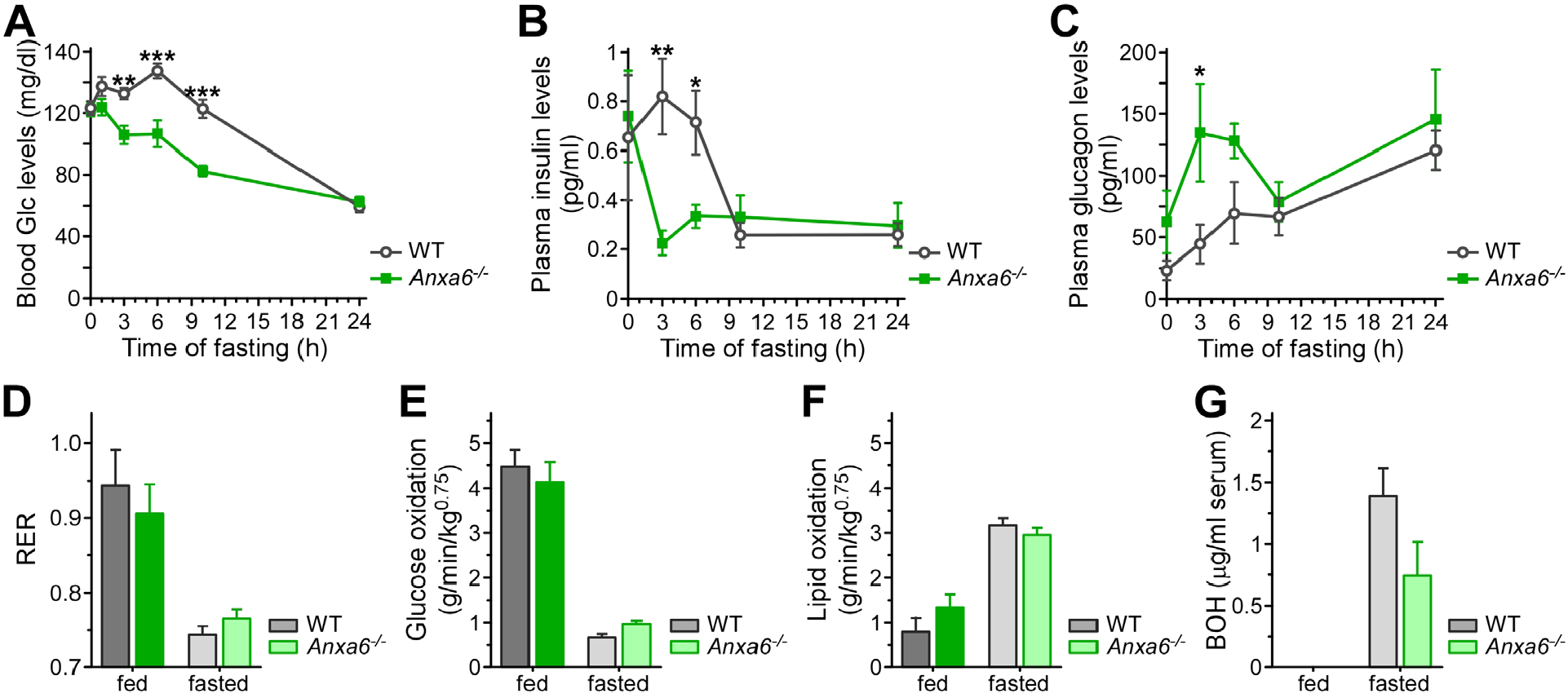
Glucose metabolism during fasting in *Anxa6*^−*/*−^ mice. (A) Blood glucose levels in WT and *Anxa6*^−*/*−^ mice (n=13 mice per group) during fasting. (B) Plasma insulin levels in WT and *Anxa6*^−*/*−^ mice (n=6-9 mice per group) during fasting. (C) Plasma glucagon levels in WT and *Anxa6*^−*/*−^ mice (n=3-6 mice per group) during fasting. (D) Respiratory exchange ratio (RER) expressed as the mean value of a fed period and 12 hour fasting period of WT and *Anxa6*^−*/*−^ mice (n=4 mice per group). (E) Glucose oxidation flux expressed as the mean value of a fed period and 12 hour fasting period of WT and *Anxa6*^−*/*−^ mice (n=4 mice per group). (F) Lipid oxidation flux expressed as the mean value of a fed period and 12 hour fasting period of WT and *Anxa6*^−*/*−^ mice (n=4 mice per group). (G) Blood β-hydroxybutyrate (BOH, ketone body) levels of WT and *Anxa6*^−*/*−^ mice before and after 12 hours fasting (n=5 mice per group).

Altogether, these results demonstrate that *Anxa6*^−*/*−^ mice were unable to maintain blood glucose levels under fasting conditions, manifested as significant hypoglycaemia immediately after food deprivation. This hypoglycaemia was produced even when glycogen stores were mobilized in the liver to secrete glucose. Interestingly, the consequent energy deficiency was alleviated using alternative energy sources such as lipid oxidation. This pointed at ANXA6 deficiency to trigger stress-induced hypoglycaemia due to deficiencies in the utilization of alternative sources for glucose production, such as GNG. In line with this hypothesis, glucagon levels were highly elevated after 0-6 hours of fasting in *Anxa6*^−*/*−^ mice (Figure 2C), supporting an increased ability to rapidly degrade glycogen and suggesting deficiencies in GNG.

### ANXA6 is essential for alanine-dependent hepatic gluconeogenesis

Besides the induction of glycogenolysis in response to prolonged hypoglycaemia, the liver also upregulates *de novo* glucose production from the utilization of non-carbohydrate carbon substrates such as pyruvate, glycerol, glutamine, and alanine. We therefore first analysed the expression of glucagon-inducible key GNG genes such as phosphoenolpyruvate carboxykinase (PEPCK, *Pck1*), glucose-6-phosphatase (*G6pc2*), and fructose-1,6-bisphosphatase (*Fbp1*) [31] in WT and *Anxa6*^−*/*−^ during the 24 hours fasting period. These data showed a trend, although not significant, towards upregulation of these genes (0-6 hours), implicating elevated GNG during these initial stages of fasting in *Anxa6*^−*/*−^ mice (Figure 3A-C). To analyse whether hepatic GNG was altered in the absence of ANXA6, we next quantified circulating glucose levels during *in vivo* tolerance tests providing pyruvate, glycerol, lactate, glutamine or alanine as substrates. No significant differences were observed in WT and *Anxa6*^−*/*−^ mice for glucose production and secretion when pyruvate and glycerol were provided as substrate (Figures 3D-E, AUC in Figure S3). In striking contrast, *Anxa6*^−*/*−^ mice showed a significantly reduced production and secretion of glucose 60 min after intraperitoneal injection of glutamine (Figures 3F, AUC in Figure S3). The lower glucose production and secretion in ANXA6-deficient animals was more prominent when lactate was provided as substrate (Figures 3G, AUC in Figure S3). Strikingly, *Anxa6*^−*/*−^ mice were unable to synthesized glucose when supplemented with alanine (Figure 3H, AUC in Figure S3), suggesting a hindrance in the glucose-alanine cycle (Cahill cycle).

**Figure 3.**
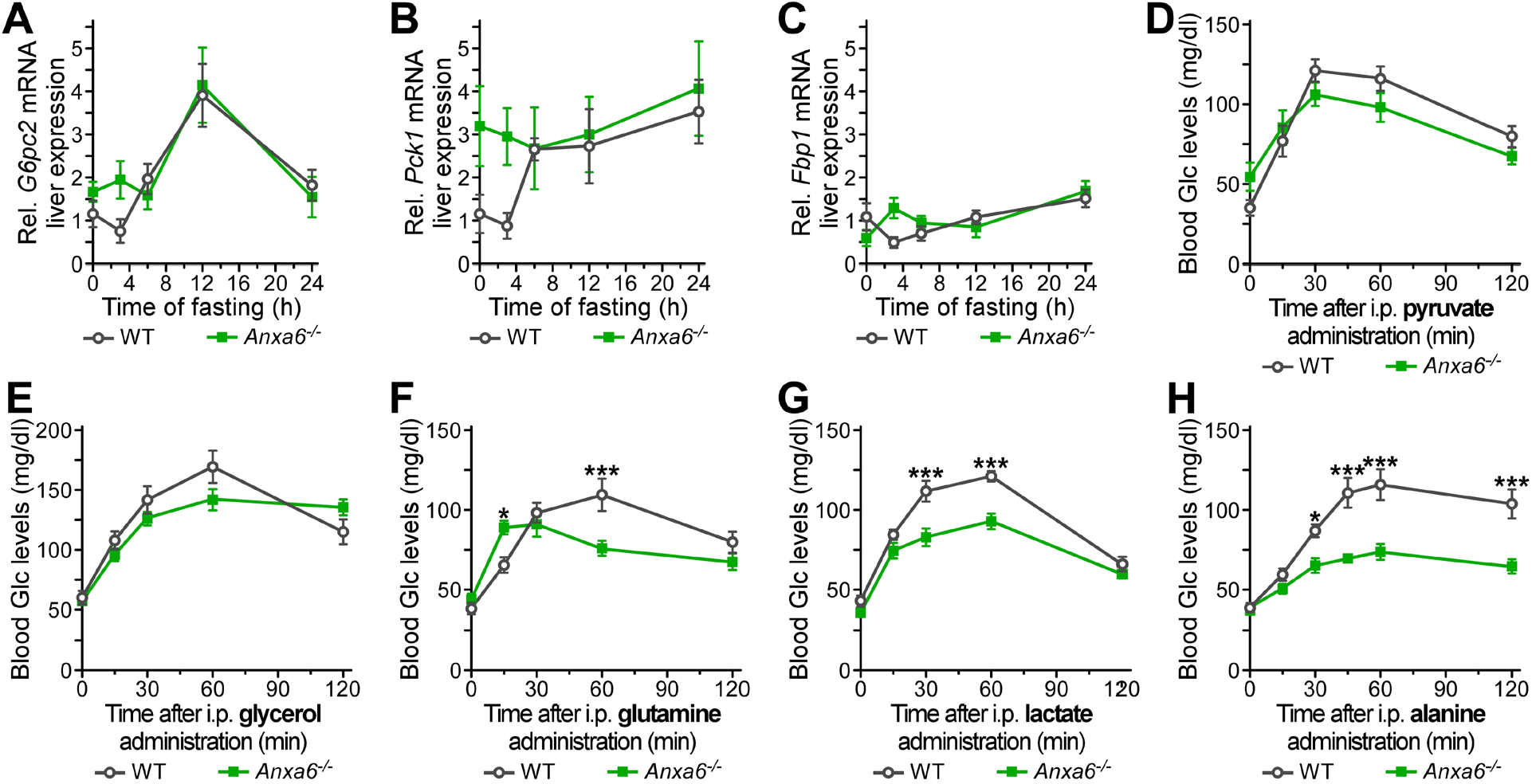
Hepatic gluconeogenic impairment in *Anxa6*^−*/*−^ mice. (A) Relative liver mRNA expression levels of glucose-6-phosphatase (*G6pc2*) during fasting in WT and *Anxa6*^−*/*−^ mice liver (n=5 mice per group and time point). (B) Relative liver mRNA expression levels of phosphoenolpyruvate carboxykinase (*Pck1*) during fasting in WT and *Anxa6*^−*/*−^ mice liver (n=5 mice per group and time point). (C) Relative liver mRNA expression levels of fructose-1,6-bisphosphatase (*Fbp1*) during fasting in WT and *Anxa6*^−*/*−^ mice liver (n=5 mice per group and time point). (D) Pyruvate tolerance test of WT and *Anxa6*^−*/*−^ mice (n=6 mice per group) after 24 hours fasting administrating i.p. 2 g/kg of sodium pyruvate. (E) Glycerol tolerance test of WT and *Anxa6*^−*/*−^ mice (n=6 mice per group) after 24 hours fasting administrating i.p. 2 g/kg of glycerol. (F) Glutamine tolerance test of WT and *Anxa6*^−*/*−^ mice (n=11 mice per group) after 24 hours fasting administrating i.p. 2 g/kg of L-glutamine. (G) Lactate tolerance test of WT and *Anxa6*^−*/*−^ mice (n=11 mice pre group) after 24 hours fasting administrating i.p. 2 g/kg of lactate. (H) Alanine tolerance test of WT and *Anxa6*^−*/*−^ mice (n=12 mice pre group) after 24 hours fasting administrating i.p. 2 g/kg of L-alanine.

**Figure S3.**
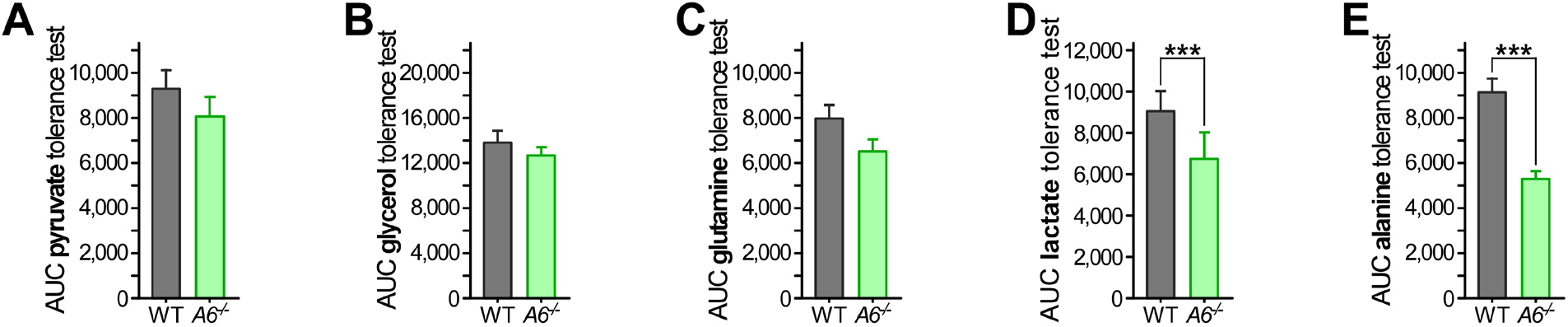
(A) Area under the curve (AUC) from pyruvate tolerance test of WT and *Anxa6*^−/−^ mice (n=6 mice per group) after 24 hours fasting administrating i.p. 2 g/kg of pyruvate. (B) Area under the curve (AUC) from glycerol tolerance test of WT and *Anxa6*^−/−^ mice (n=6 mice per group) after 24 hours fasting administrating i.p. 2 g/kg of glycerol. (C) Area under the curve (AUC) from glutamine tolerance test of WT and *Anxa6*^−/−^ mice (n=11 mice per group) after 24 hours fasting administrating i.p. 2 g/kg of glutamine. (D) Area under the curve (AUC) from lactate tolerance test of WT and *Anxa6*^−/−^ mice (n=11 mice per group) after 24 hours fasting administrating i.p. 2 g/kg of lactate. (E) Area under the curve (AUC) from alanine tolerance test of WT and *Anxa6*^−/−^ mice (n=12 mice per group) after 24 hours fasting administrating i.p. 2 g/kg of alanine.

Taken together, these results demonstrate the limited capacity of *Anxa6*^−*/*−^ mice to produce and secrete glucose from glutamine and lactate, and their complete inability to use alanine as a substrate for hepatic GNG *in vivo*.

### ANXA6 deficiency do not affect alanine metabolization capability of the liver

Based on the data shown above, the failure of *Anxa6*^−*/*−^ mice to maintain blood glucose levels during fasting was not due to a dysfunction of the glucose and insulin regulatory axis, but the inability to produce glucose mainly from alanine, the key gluconeogenic substrate during fasting. To substantiate these findings, we next compared amino acid levels in serum and liver extracts from fed and 24 hours fasted WT and *Anxa6*^−*/*−^ mice (relative representation of amino acid levels in Figure 4A and 4B, complete amino acid profile in Supporting Table 3 and 4). As expected, alanine plasma levels (detailed in Figure 4C) significantly decreased upon fasting in WT mice. In contrast, plasma alanine concentrations in *Anxa6*^−*/*−^ mice remained unaltered after fasting, further supporting an inability of *Anxa6*^−*/*−^ mice to utilize alanine as GNG substrate or reduced alanine muscle secretion. Furthermore, liver alanine levels were comparable in both strains of mice (Figure 4D). Yet, elevation of liver glutamate levels after fasting was much more prominent in WT compared to *Anxa6*^−*/*−^ animals (Figure 4E), indicating a higher metabolic flux from alanine to pyruvate in WT mice. Urine urea levels, an additional indicator of alanine and glutamine deamination, exhibited an increase in WT mice; however, urine urea amount remained constant in fasted *Anxa6*^−*/*−^ mice (Figure 4F). The determination of all other amino acid levels in fed and fasted WT and *Anxa6*^−*/*−^ mice in plasma or liver did not reveal any other major change possibly being responsible for a defect in hepatic GNG in fasted *Anxa6*^−*/*−^ animals. Thus, based on these analyses, impaired ability to drive hepatic GNG via alanine appears a likely factor for the development of hypoglycaemia in fasted *Anxa6*^−*/*−^ mice.

**Figure 4.**
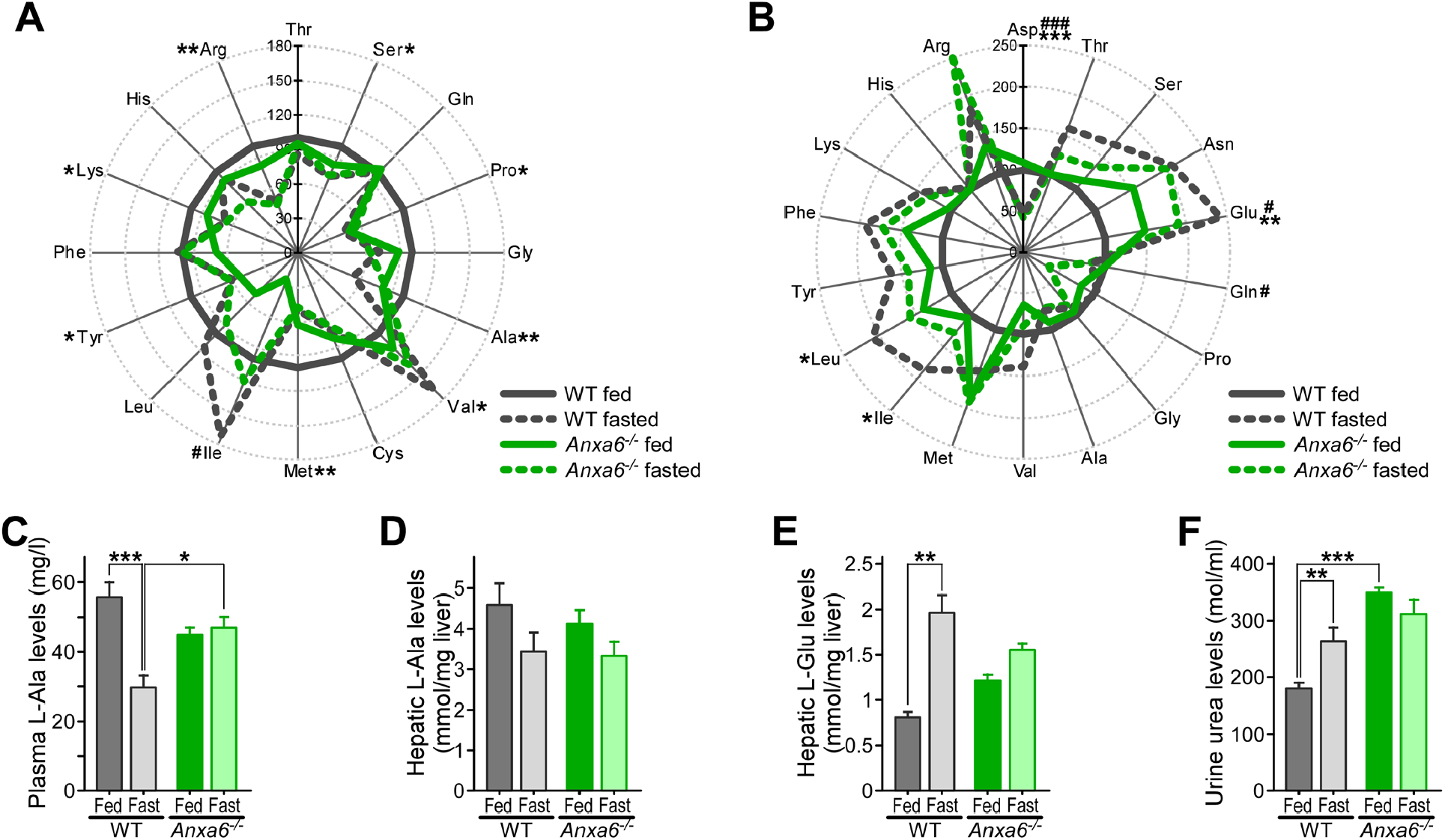
ANXA6 deficiency do not affect alanine metabolization capability of the mice liver. (A) Spider diagram representation of relative plasma threonine (L-Thr), serine (L-Ser), glutamine (L-Gln), proline (L-Pro), glycine (L-Gly), alanine (L-Ala), valine (L-Val), cysteine (L-Cys), methionine (L-Met), isoleucine (L-Ile), leucine (L-Leu), tyrosine (L-Tyr), phenylalanine (L-Phe), lysine (L-Lys), histidine (L-His) and arginine (L-Arg) levels of WT and *Anxa6*^−*/*−^ mice fed and fasted for 24 hours (n=4 mice per group). (B) Spider diagram representation of relative hepatic aspartic acid (L-Asp), threonine (L-Thr), serine (L-Ser), asparagine (L-Asn), glutamic acid (L-Glu), glutamine (L-Gln), proline (L-Pro), glycine (L-Gly), alanine (L-Ala), valine (L-Val), methionine (L-Met), isoleucine (L-Ile), leucine (L-Leu), tyrosine (L-Tyr), phenylalanine (L-Phe), lysine (L-Lys), histidine (L-His) and arginine (L-Arg) levels of WT and *Anxa6*^−*/*−^ mice fed and fasted for 24 hours (n=4 mice per group). (C) Plasma alanine levels in WT and *Anxa6*^−*/*−^ mice fed and fasted for 24 hours (n=4 each group). (D) Hepatic alanine levels in WT and *Anxa6*^−*/*−^ mice fed and fasted for 24 hours (n=4 each group). (E) Hepatic glutamic acid levels in WT and *Anxa6*^−*/*−^ mice fed and fasted for 24 hours (n=4 each group). (F) Urine urea levels in WT and *Anxa6*^−*/*−^ mice fed and fasted for 24 hours (n=4 each group).

To further dissect and identify de-regulated mechanisms of the alanine-glucose cycle in ANXA6-deficient hepatocytes, we analysed the expression of hepatic enzymes converting alanine into pyruvate: alanine aminotransferase (ALAT) 1 and 2 (*Gpt1/2*). In response to fasting, reduced *Gpt2* levels were observed in *Anxa6*^−*/*−^ livers (Figure 5A-B), yet ALAT activity was similar in liver extracts from WT and *Anxa6*^−*/*−^ mice (Figure 5C), indicating that reduced *Gpt2* mRNA levels did not significantly impact on enzyme activity. In addition, hepatic lactate dehydrogenase (*Ldha*) was slightly reduced in *Anxa6*^−*/*−^ mice after 24 hours fasting (Figure 5D). Hence, at least at the mRNA level, hepatic expression of enzymes metabolizing alanine into pyruvate (Figure 5A-C) and lactate into pyruvate (Figure 5D) were slightly reduced in the absence of ANXA6 after prolonged fasting, although their enzyme activity nor those enzymes converting pyruvate into glucose (Figure 3) appeared significantly affected.

**Figure 5.**
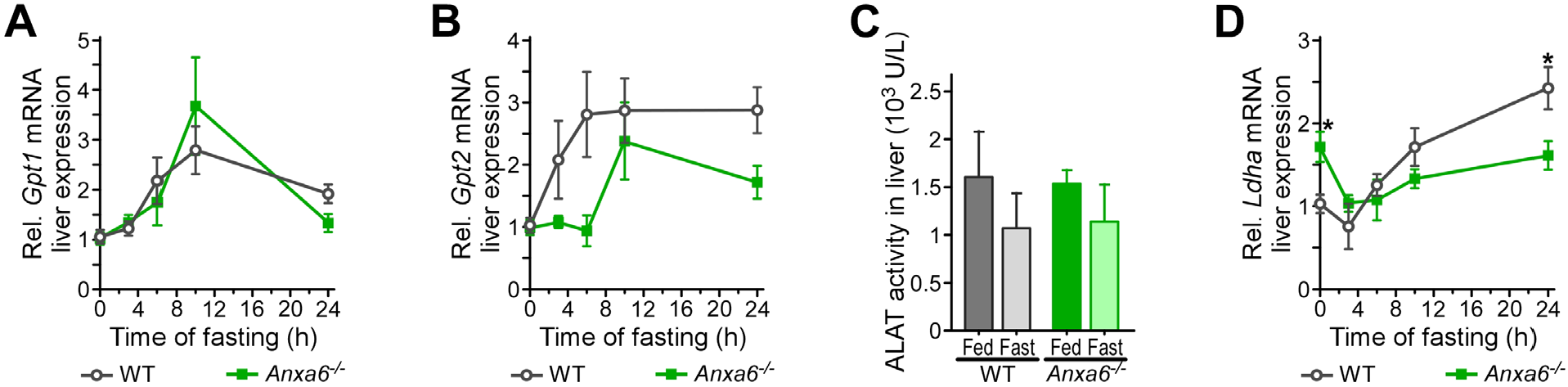
Hepatic alanine metabolization capacity in *Anxa6*^−*/*−^ mice. (A) Relative liver mRNA expression levels of alanine aminotransferase 1 (*Gpt1*) during fasting in WT and *Anxa6*^−*/*−^ mice liver (n=5 per group and time point). (B) Relative liver mRNA expression levels of alanine aminotransferase 2 (*Gpt2*) during fasting in WT and *Anxa6*^−*/*−^ mice liver (n=5 per group and time point). (C) Alanine aminotransferase activity in WT and *Anxa6*^−*/*−^ liver during fasting (n=5 per group and time point). (D) Relative liver mRNA expression levels of lactate dehydrogenase (*Ldha2*) during fasting in WT and *Anxa6*^−*/*−^ mice liver (n=5 per group and time point).

### Hepatic alanine transporters during fasting in *Anxa6*^−*/*−^ mice

Similar to the lack of blood alanine clearance upon fasting in *Anxa6*^−*/*−^ mice (Figure 4C), we previously demonstrated *Anxa6*^−*/*−^ mice unable to use blood alanine for GNG after partial hepatectomy. This correlated with a failure to internalize alanine in *Anxa6*^−*/*−^ hepatocytes [10], pointing at compromised hepatic uptake of alanine as a potentially underlying mechanism for the impaired alanine-promoted GNG of fasted *Anxa6*^−*/*−^ mice.

We therefore first analysed mRNA levels of SNAT2 and 4 (*Slc38a2* and *Slc38a4*) and protein levels of SNAT4 (Figure 6A-D). Elevation of hepatic SNAT2 mRNA levels was delayed in *Anxa6*^−*/*−^ mice during fasting, while SNAT4 mRNA expression remained constant over the fasting period (Figure 6A-B). Furthermore, elevation of SNAT4 protein levels after 24 hours in WT mice was not observed in ANXA6-deficient animals (Figure 6C, quantification in 6D). Together, these findings may indicate reduced amounts of SNATs being available for alanine uptake in fasted *Anxa6*^−*/*−^ livers.

**Figure 6.**
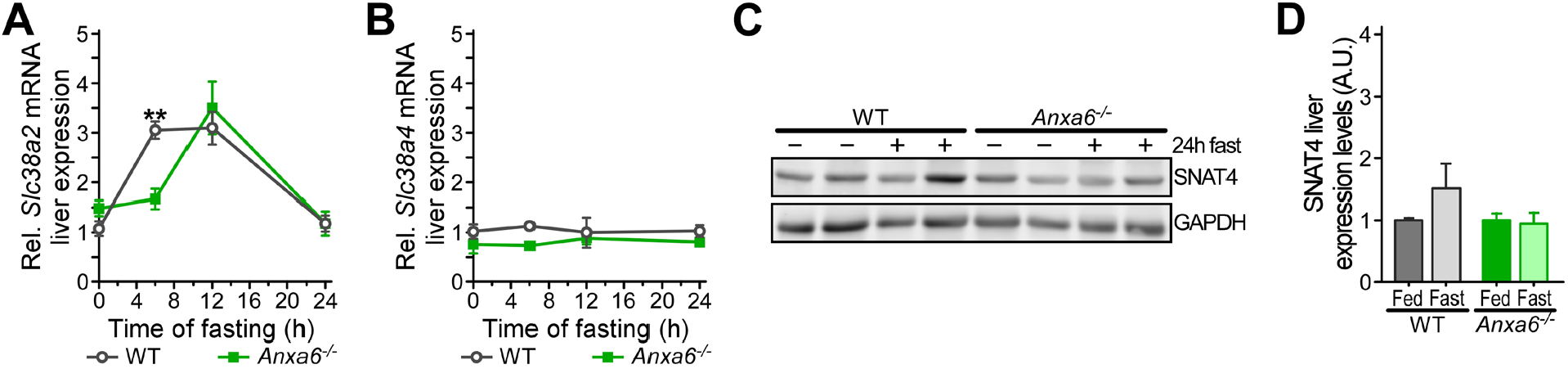
Hepatic alanine transporters during fasting in *Anxa6*^−*/*−^ mice. (A) Relative liver mRNA expression levels of SNAT2 (*Slc38a2*) after 24 hours fasting in WT and *Anxa6*^−*/*−^ mice liver (n=5 per group and time point). (B) Relative liver mRNA expression levels of SNAT4 (*Slc38a4*) after 24 hours fasting in WT and *Anxa6*^−*/*−^ mice liver (n=5 per group and time point). (C) Relative expression of SNAT4 amino acid transporter after 24 hours fasting in WT and *Anxa6*^−*/*−^ mice liver (n=2 per group and time point). (D) Relative quantification of SNAT4 amino acid transporter expression after 24 hours fasting in WT and *Anxa6*^−*/*−^ mice liver (n=4 mice per group and time point).

Altogether, *Anxa6*^−*/*−^ mice showed an energetic imbalance during fasting that induced a fast drop in blood glucose levels and a faster switch from glycolytic to lipolytic metabolism without any indication of the development of a diabetic phenotype. In *Anxa6*^−*/*−^ mice, the inability to maintain glycolytic metabolism in the initial stages of fasting was due to an impairment in alanine-dependent GNG.

## DISCUSSION

The objective of this study was to examine the function of ANXA6 in maintaining glucose homeostasis under physiological conditions that reflected the fed and fasted state. *Anxa6*^−*/*−^ mice exhibited glucose absorption and insulin response that was comparable to WT mice during the absorptive state. These findings were anticipated following the previous analysis of glucose metabolism after partial hepatectomy or upon a prolonged feeding of a high-fat diet [10, 25]. However, our findings indicate that ANXA6 plays a pivotal role in maintaining blood glucose levels during fasting, despite the apparent normal glucose metabolism observed under feeding conditions. The metabolic response analysis over a 24-hour period using indirect calorimetry demonstrated that both WT and *Anxa6*^−*/*−^ mice initiated lipid oxidation-dependent energy production during the initial hours of the quiescent dark period. Though, the *Anxa6*^−*/*−^ mice were unable to induce the lipid-to-carbohydrate catabolic switch at the end of the dark period in comparison to their control littermates. This was probably due to the inability of the *Anxa6*^−*/*−^ mice to activate the hepatic gluconeogenic pathway, as observed during liver regeneration following partial hepatectomy [10]. In accordance, the insulin-sensitive control of systemic glucose levels, glucose absorption and secretion were not negatively affected in *Anxa6*^−*/*−^ mice during regular feeding conditions.

Interestingly, our data demonstrated a notable decrease in blood glucose levels during the first stages of fasting in *Anxa6*^−*/*−^ mice, in comparison to WT mice. This initially suggested an impairment of hepatic GNG or glycogen mobilization. However, the analysis of the expression of gluconeogenic key regulatory genes (*G6pc2, Pck1* and *Fbp1*) revealed elevated levels of PEPCK expression in the liver during the basal fed state in *Anxa6*^−*/*−^ mice, indicating its predisposition to GNG activation even during the fed state. A comparable phenotype has been observed in the liver glycogen synthase knockout (*Gys2*^−*/*−^) mice, which exhibited elevated hepatic *Pck1* gene expression and PEPCK activity during the fed state [32]. Nevertheless, *Gys2*^−*/*−^ mice displayed a 95% reduction in liver glycogen content during the fed state and exhibited insulin resistance [32, 33], which was not observed in *Anxa6*^−*/*−^ mice. The robust fasting-induced hypoglycaemia observed in *Anxa6*^−*/*−^ mice persisted despite the efficient mobilisation of elevated hepatic glycogen reserves at elevated degradation rates from the onset of the fasting state in these animals. This phenomenon correlated with a rapid decline in blood insulin levels and a rapid increase in blood glucagon levels. Taken together these data suggest a very sensitive and elevated reaction capacity of *Anxa6*^−*/*−^ hepatocytes facing a carbohydrate-dependent energetic deficit, inducing a fast switch to a lipid-catabolic metabolism.

The pronounced hypoglycaemia during the fasting state observed in *Anxa6*^−*/*−^ mice correlated with its inability to sustain glucose production via GNG, particularly from alanine, the major gluconeogenic substrate in fasted WT mice [34]. The pyruvate and glycerol tolerance tests indicated that there was no significant difference in blood glucose secretion between WT and *Anxa6*^−*/*−^ mice. These results suggest that the inability to produce *de novo* glucose observed in *Anxa6*^−*/*−^ mice was not due to a biochemical defect, but rather a limitation in the availability of GNG substrates during fasting. In accordance, the urea cycle, which is essential for the detoxification of ammonium ions released during amino acid catabolism [35], appeared fully functional in *Anxa6*^−*/*−^ mice as indicated by high urine urea levels during both fed and fasting states. Notably, data suggested that the urea cycle was continuously activated in *Anxa6*^−*/*−^ mice due to a defective amino acid metabolism and/or turnover, which correlated with high urine production and water intake of *Anxa6*^−*/*−^ mice. *Anxa6*^−*/*−^ mice also showed lower glucose production via GNG from lactate but not pyruvate, which may relate to the lower *Ldha* expression levels observed in these *Anxa6*^−*/*−^ mice during fasting.

The availability of gluconeogenic substrates in the liver is a critical determinant of the GNG flux rates [2]. Alanine availability for the liver is not affected in *Anxa6*^−*/*−^ compared to WT mice, which showed lower but not significantly reduced levels of blood alanine during the fed state. As anticipated, 24-hour fasting resulted in a reduction of blood alanine levels in WT mice, indicating the presence of a robust GNG-dependent alanine uptake mechanism. The slight reduction in hepatic alanine levels and the increase in urine urea levels were also found to correlate with this alanine-dependent GNG in WT mice. Interestingly, *Anxa6*^−*/*−^ mice exhibited comparable blood alanine levels during both fed and fasting states, despite the absence of discernible alterations in hepatic ALAT activity. These findings indicate a consistent alanine metabolic capacity, yet an inoperative alanine uptake in hepatocytes during fasting in *Anxa6*^−*/*−^ mice, which is analogous to our previous results observed in *Anxa6*^−*/*−^ mice during liver regeneration [10].

Earlier research in *Anxa6*^−*/*−^ mice demonstrated that SNAT4 recycling to the sinusoidal plasma membrane after partial hepatectomy in *Anxa6*^−*/*−^ mice was impaired, which correlated with compromised alanine uptake and *de novo* glucose production [10]. Concurrently, Curnock and colleagues demonstrated that the nutrient-sensing and adaptive delivery of SNAT2 to the plasma membrane depends on the retromer complex, which prevents SNAT2 degradation in lysosomes [13]. On the other hand, the promoter of the retromer complex subunits *VPS35, VPS26A* and *SNX27* genes contain TFEB/TFE3-responsive elements, which regulate its expression in response to amino acid starvation or selective glutamine depletion [13]. Then, the nutrient-dependent expression of retromer proteins has been shown to modulate the retromer-mediated retrieval and recycling of amino acid transporter proteins. Intriguingly, SNAT2 gene expression is upregulated during fasting, suggesting a further level of nutrient-dependent regulation of this amino acid transporter. The recruitment and assembly of retromer components requires active RAB7 (RAB7-GTP) [14]. In contrast, upregulation of the RAB7-GAP TBC1D5 inhibits retromer complex formation through downregulating RAB7 activity [15]. In addition, ANXA6 has been demonstrated to directly interact with the RAB7-GAP TBC1D15 to regulate RAB7-GTP levels, with consequences for intracellular cholesterol trafficking [22]. These studies may point at a yet unknown link between ANXA6 and RAB7 for the regulation of SNAT localization in a nutrient-sensing manner.

Altogether, our findings identify that ANXA6 deficiency causes an inability to maintain glycolytic metabolism under fasting conditions due to impaired alanine-dependent GNG.

## Abbreviations

ALAT: alanine aminotransferase
ANXA6: Annexin A6
AUC: area under the curve
BOH: beta-hydroxybutyrate
FBP: fructose-1,6-bisphosphatase
G6Pase: glucose-6-phosphatase
GAP: GTPase activating protein
GNG: gluconeogenesis; i.p., intraperitoneal
LDH: lactate dehydrogenase
PEPCK: phosphoenolpyruvate carboxykinase
RER: respiratory exchange ratio
SNAT: sodium-coupled neutral amino acid transporter
T2DM: type 2 diabetes mellitus
WT: wild type

