## Supplementary material for "Fasting-induced hepatic gluconeogenesis is compromised in *Anxa6^−/−^* mice": Table 3 and 4

**Table 3.** Plasma amino acid levels (mg/l) of WT and *Anxa6*$^{-/-}$ mice fed and fasted for 24 hours.

| | WT_1 fed | WT_2 fed | WT_3 fed | WT_4 fed | WT_1 fasted | WT_2 fasted | WT_3 fasted | WT_4 fasted | A6ko_1 fed | A6ko_2 fed | A6ko_3 fed | A6ko_4 fed | A6ko_1 fasted | A6ko_2 fasted | A6ko_3 fasted | A6ko_4 fasted |
|---|---|---|---|---|---|---|---|---|---|---|---|---|---|---|---|---|
| Ala | 68.31 | 51.54 | 50.37 | 52.69 | 34.99 | 36.06 | 26.33 | 21.69 | 49.41 | 40.15 | 47.08 | 43.40 | 41.59 | 46.18 | n.a. | 52.85 |
| Thr | 33.06 | 25.28 | 25.40 | 27.79 | 21.24 | 27.90 | 23.35 | 22.93 | 24.36 | 19.30 | 24.63 | 37.65 | 31.61 | 30.23 | 15.15 | 29.01 |
| Ser | 21.95 | 16.56 | 16.47 | 16.62 | 12.59 | 15.83 | 10.97 | 11.81 | 15.39 | 13.63 | 13.09 | 17.55 | 14.50 | 15.87 | 7.41 | 14.10 |
| Gln | 66.00 | 64.81 | 91.85 | 83.07 | 61.86 | 76.65 | 86.85 | 78.78 | 84.26 | 76.02 | 94.42 | 111.64 | 66.39 | 78.45 | 36.07 | 109.05 |
| Pro | 16.03 | 20.21 | 15.00 | 11.20 | 9.56 | 8.65 | 9.29 | 0.00 | 11.62 | 7.58 | 11.35 | 0.00 | 9.39 | 13.17 | 0.00 | 11.42 |
| Gly | 29.07 | 26.91 | 23.96 | 18.44 | 18.14 | 18.74 | 18.03 | 15.80 | 22.99 | 17.49 | 19.05 | 27.27 | 17.18 | 17.64 | 9.80 | 17.81 |
| Val | 33.27 | 35.47 | 33.29 | 27.67 | 23.58 | 14.05 | 21.25 | 31.87 | 29.60 | 32.58 | 14.86 | 28.80 | 29.60 | 32.58 | 14.86 | 28.80 |
| Cys | 12.90 | 11.03 | 19.44 | 13.89 | 8.92 | 9.61 | 10.24 | 12.62 | 11.99 | 11.94 | 12.71 | 9.49 | 9.52 | 11.70 | 6.99 | 11.01 |
| Met | 15.17 | 14.06 | 16.71 | 9.95 | 4.72 | 8.61 | 8.71 | 6.00 | 13.07 | 11.03 | 10.85 | 0.00 | 12.00 | 9.39 | 0.00 | 4.79 |
| Ile | 14.99 | 0.00 | 5.69 | 10.85 | 12.87 | 14.25 | 11.39 | 16.14 | 0.00 | 0.00 | 7.85 | 0.00 | 10.43 | 12.95 | 5.84 | 9.05 |
| Leu | 35.01 | 24.56 | 19.63 | 20.44 | 31.38 | 32.47 | 24.94 | 25.57 | 17.33 | 16.00 | 16.25 | 0.00 | 25.67 | 26.04 | 10.11 | 24.16 |
| Tyr | 31.84 | 31.80 | 17.32 | 28.21 | 17.27 | 17.27 | 16.60 | 15.66 | 18.30 | 21.58 | 18.79 | 0.00 | 15.65 | 20.29 | 8.14 | 23.85 |
| Phe | 18.72 | 16.02 | 10.54 | 11.68 | 14.24 | 16.57 | 11.61 | 16.87 | 12.95 | 13.69 | 13.55 | 0.00 | 18.24 | 17.38 | 8.36 | 13.18 |
| Lys | 71.68 | 55.41 | 45.36 | 52.79 | 41.18 | 36.17 | 38.38 | 36.02 | 50.08 | 50.36 | 48.39 | 40.83 | 44.49 | 37.50 | 26.56 | 42.18 |
| Arg | 34.18 | 25.42 | 22.68 | 28.48 | 11.97 | 16.04 | 13.12 | 11.60 | 21.46 | 20.26 | 24.95 | 24.21 | 18.31 | 14.11 | 0.00 | 17.09 |
| His | 14.60 | 13.82 | 13.00 | 9.74 | 12.26 | 11.34 | 10.57 | 13.17 | 10.78 | 10.10 | 9.81 | 15.13 | 10.23 | 9.87 | 4.10 | 7.93 |

**Table 4**. Hepatic amino acid levels (µmol/g) of WT and *Anxa6*$^{-/-}$ mice fed and fasted for 24 hours.

| | WT_1 fed | WT_2 fed | WT_3 fed | WT_4 fed | WT_1 fasted | WT_2 fasted | WT_3 fasted | WT_4 fasted | A6ko_1 fed | A6ko_2 fed | A6ko_3 fed | A6ko_4 fed | A6ko_1 fasted | A6ko_2 fasted | A6ko_3 fasted | A6ko_4 fasted |
|---|---|---|---|---|---|---|---|---|---|---|---|---|---|---|---|---|
| Asp | 3527.76 | 5189.25 | 4960.31 | 5032.15 | 2236.70 | 2149.51 | 2360.62 | 1857.01 | 4856.23 | 5355.09 | 5352.55 | 4739.71 | 1676.29 | 2003.01 | 1748.42 | 2007.26 |
| Thr | 563.87 | 748.46 | 416.90 | 248.22 | 1042.81 | 1034.45 | 436.90 | 623.15 | 474.66 | 453.56 | 561.76 | 497.44 | 791.02 | 802.17 | 436.12 | 430.31 |
| Ser | 893.82 | 1064.66 | 763.57 | 405.55 | 1862.69 | 1690.24 | 620.62 | 1139.29 | 775.47 | 845.05 | 1024.76 | 907.43 | 1401.77 | 1340.47 | 758.04 | 727.93 |
| Asn | 270.44 | 208.57 | 475.66 | n.a. | 688.69 | 607.53 | 344.81 | 364.29 | 353.66 | 358.58 | 400.69 | 354.81 | 729.75 | 544.15 | 349.05 | 318.99 |
| Glu | 750.96 | 994.74 | 788.68 | 701.28 | 2223.24 | 2360.95 | 1571.32 | 1709.13 | 1133.33 | 1117.02 | 1388.58 | 1229.60 | 1436.97 | 1760.37 | 1541.48 | 1478.07 |
| Gln | 4524.48 | 3397.29 | 3672.56 | 3912.52 | 3144.65 | 4033.05 | 2852.71 | 2990.61 | 4307.45 | 4094.56 | 3787.35 | 3353.72 | 3029.46 | 3405.52 | 3080.76 | 3372.49 |
| Pro | 165.12 | 334.60 | 91.32 | 166.29 | 201.07 | 292.05 | 196.12 | 93.65 | 257.45 | n.a. | 190.37 | 168.58 | n.a. | n.a. | 95.27 | 179.05 |
| Gly | 2886.83 | 3563.15 | 3125.58 | 2204.13 | 2933.33 | 3056.76 | 2071.47 | 2428.04 | 2990.24 | 3171.55 | 2729.44 | 2416.93 | 2489.99 | 2718.23 | 2289.12 | 2307.12 |
| Ala | 3347.65 | 5902.22 | 4238.30 | 4869.84 | 4377.98 | 3570.43 | 3714.26 | 2081.09 | 5062.87 | 3640.59 | 4163.14 | 3686.48 | 2823.42 | 3668.90 | 2708.68 | 4158.52 |
| Val | 519.34 | 577.74 | 139.38 | 220.63 | 359.48 | 811.58 | 335.50 | 530.29 | 226.15 | n.a. | 372.22 | 329.60 | 331.22 | 418.06 | 243.06 | 293.86 |
| Met | 250.89 | 334.08 | 233.49 | n.a. | 690.83 | 619.11 | 279.54 | 371.69 | 273.31 | 247.00 | 340.44 | 301.46 | 517.53 | 469.90 | 297.48 | 357.96 |
| Ile | 242.76 | 199.76 | 126.51 | 125.75 | 388.99 | 424.55 | 215.50 | 274.74 | 136.04 | 137.52 | 239.34 | 211.94 | 235.35 | 308.86 | 159.94 | 193.02 |
| Leu | 1053.52 | 1614.93 | 973.80 | 388.91 | 2692.05 | 2768.90 | 1351.01 | 1692.20 | 1176.69 | 1381.17 | 1629.71 | 1443.12 | 1961.41 | 2004.68 | 1073.34 | 1356.15 |
| Tyr | 467.39 | 530.42 | 274.73 | 184.35 | 827.68 | 672.11 | 397.83 | 459.92 | 398.10 | 427.48 | 440.30 | 389.89 | 593.52 | 711.04 | 333.75 | 410.34 |
| Phe | 458.11 | 757.79 | 475.04 | 162.45 | 1271.10 | 1160.25 | 500.00 | 664.15 | 644.99 | 678.94 | 728.47 | 645.07 | 1002.36 | 953.68 | 549.69 | 676.54 |
| Lys | 930.93 | 838.09 | 970.70 | 684.92 | 1590.67 | 1345.05 | 1129.77 | 858.20 | 771.68 | 851.19 | 1057.50 | 936.42 | 1319.59 | 1338.80 | 1049.69 | 854.33 |
| His | 799.77 | 814.30 | 775.97 | 554.05 | 962.84 | 735.29 | 662.02 | 651.59 | 658.27 | 693.45 | 822.01 | 727.89 | 893.37 | 772.58 | 638.49 | 629.75 |
| Arg | 182.10 | 156.66 | 284.96 | n.a. | 571.87 | 271.69 | 164.03 | 144.44 | 175.20 | 279.22 | 203.99 | 180.63 | 640.21 | 506.36 | 228.86 | 181.70 |
